# Dimethyl fumarate reprograms cervical cancer cells to enhance antitumor immunity by activating mtDNA-cGAS-STING pathway

**DOI:** 10.1101/2025.04.27.650830

**Authors:** Han Jiang, Liting Liu, Shan He, Shen Qu, Yifan Yang, Guijie Kang, Min Wu, Hangyu Liu, Yuwei Zhang, Zixuan Wang, Wenjing Tian, Ying Chen, Liming Wang, Qiangqiang Wang, Ting Ye, Junyan Han, Hui Wang, Yafei Huang

## Abstract

**Background:** Cervical cancer (CC) remains a significant global health challenge for women, especially in advanced stages where effective treatments are limited. Current immunotherapies, including PD-1/PD-L1 blockades and adoptive T cell therapies, show limited response rates and durability. Dimethyl fumarate (DMF), an FDA-approved drug for autoimmune diseases, has demonstrated that it might exhibit direct antitumor activity in several cancers. However, its influence on anti-tumor immunity and its function in CC remain poorly understood. This study aims to investigate the therapeutic potential of DMF in CC models and elucidate its underlying mechanisms of action.

**Methods:** CC cell lines and mouse models were treated with DMF. Transcriptional level changes in cervical cancer cells following DMF treatment were analyzed by RNA-seq. Mitochondrial DNA (mtDNA) release, and cGAS-STING activation were assessed via qPCR, immunofluorescence, immunoblotting and ELISA. CD8^+^ T cell recruitment was analyzed by flow cytometry. Combinatorial therapies (DMF + anti-PD-1/TILs) were tested in syngeneic or patient-derived xenografts (PDX) models.

**Results:** DMF treatment induces mitochondrial dysfunction in tumor cells, resulting in the release of mtDNA into the cytosol. The cytosolic mtDNA activates the cGAS-STING-TBK1 pathway and type I interferon response, leading to the secretion of CCL5 and CXCL10, thereby enhancing CD8⁺ T cell infiltration. Additionally, DMF exhibits synergistic effect with PD-1 blockade in murine CC model, and can enhance the therapeutic efficacy of adoptively transferred T cells toward CC patient-derived xenografts model.

**Conclusion:** This work elucidated that DMF reprograms CC cells to activate the mtDNA-cGAS-STING pathway, fostering a chemokine-rich microenvironment that recruits CD8^+^ T cells. Its synergy with PD-1 blockade or TIL therapy underscores its potential as an immunomodulatory adjuvant. These findings suggest that DMF holds promise as a novel immunotherapeutic strategy for improving clinical outcomes in CC.

## Introduction

Cervical cancer (CC) is one of the most common malignant tumors in women, ranking first in incidence and third in mortality among gynecological tumors worldwide, and also occupying a leading position among female malignancies (1–3). While early-stage CC is often curable through surgery, radiotherapy, and chemotherapy, effective treatments for late-stage and recurrent metastatic cases are still lacking. As a result, these patients suffer from a poor prognosis, with an overall survival of 7 to 9 months and a 5-year survival rate of 15.5% (4). Therefore, developing new therapeutic strategies is urgently needed.

Tumor immunotherapy has gained attention in CC treatment due to its promising potential (5). However, current immunotherapies still fall short of clinical expectations. Approved immune checkpoint blockades (ICBs), such as anti-PD-1/PD-L1 antibodies, show objective response rates of only 14.6%-27.8% in advanced CC patients (6–8), highlighting issues with immunogenicity and durability. Meanwhile, as a promising therapeutic modality by replenishing tumor-specific T cells, T cell receptor-gene engineered T cell (TCR-T) therapy is limited by antigen selection and MHC restriction, benefiting only a small number of patients (9–11). This obstacle can be circumvented by tumor-infiltrating lymphocyte (TIL) therapy through targeting multiple tumor antigens; however, the efficacy of TIL varies due to patient heterogeneity (12, 13). Therefore, improving the efficacy of current immunotherapies is of paramount importance for treating CC patients.

Dimethyl fumarate (DMF) is an FDA-approved clinical drug for the treatment of autoimmune diseases such as multiple sclerosis and psoriasis (14, 15). It is mainly administered orally and is rapidly converted into its primary active metabolite, monomethyl fumarate (MMF), under the action of esterases in the body. Both DMF and MMF exhibit excellent cell permeability (absorption capacity) (16, 17). Interestingly, while DMF primarily exerts anti-inflammatory effects in autoimmune diseases, it mediates antitumor effects in cancer. In preclinical models, DMF treatment has been shown to cause regression and/or reduced metastasis in various types of tumors, such as melanoma (18, 19), colorectal cancer (20, 21), breast cancer (22, 23), and lung cancer (24). However, despite extensive research supporting the antitumor potential of DMF, its antitumor mechanism, particularly its impact on antitumor immunity, remains unclear.

Here, we demonstrated that DMF is capable of reprogramming CC cells to promote anti-tumor immunity through activating the mtDNA-cGAS-STING pathway. DMF treatment induces mitochondrial dysfunction in CC cells, leading to the release of mitochondrial DNA (mtDNA) into the cytoplasm, which activates the cGAS-STING pathway and type I interferon response, and subsequently induces the secretion of CCL5 and CXCL10, thereby promoting CD8^+^ T cell recruitment. Importantly, combinatorial therapy with DMF and PD-1 blockade is superior to monotherapies, suggesting a synergistic effect. Finally, DMF is able to potentiate the therapeutic efficacy of adoptively transferred TILs in the human patient-derived xenografts (PDX) model of CC, suggesting its potential in clinical application.

## Materials and Methods

### Cell lines

The murine tumor cell line TC-1 and human tumor cell line SiHa were purchased from the American Type Culture collection (ATCC, USA). All cell lines were cultured in DMEM supplemented with 10% fetal bovine serum (FBS) at 37℃ in 5% CO_2_ and routinely tested for mycoplasma. *Sting* KO TC-1 cells and *Ccl5*&*Cxcl10* KO TC-1 cells were generated using CRISPR plasmid vectors for the mouse *Sting* gene (VectorBuilder, VB900088-3817jne) and *Ccl5*&*Cxcl10* genes (VectorBuilder, VB240830-1104jfz), respectively. In brief, the CRISPR plasmid vectors were transfected into target cells using lipofectamine (Invitrogen, L3000015). At 48 h post-transfection, puromycin (5 μg/mL) was added to the culture medium to initiate selection, which continued for 5 days to eliminate non-transfected cells and enrich populations stably expressing the resistance gene. Surviving cells were then seeded by limiting dilution in 96-well plates to establish monoclonal colonies.

### Mice

C57BL/6J and BALB/c nude mice were purchased from the Beijing Vital River Laboratory (Beijing, China). NOD/ShiLtJGpt-*Prkdc*^em26Cd52^*Il2rg*^em26Cd22^/Gpt (NCG) mice were purchased from GemPharmatech (Nanjing, China). All mice were kept under specific-pathogen-free conditions with controlled temperature (21 °C–23 °C), humidity (30–60%) and light cycle (12-h light/dark) at the Animal Care and Use Committee of Tongji Medical College. Six- to eight-week-old female mice were used in all experiments.

### Antibodies and reagents

For flow cytometry, Fixable viability stain (564406) was purchased from BD; anti-mouse Fc Blocker (S17011E), CD45 (30-F11), CD3ε (145-2C11), CD4 (GK1.5), CD8α (53-6.7), IFN-γ (XMG1.2), TNF-α (MP6-XT22), Granzyme B (QA16A02), perforin (S16009A); anti-human Fc Blocker (422305), CD45 (2D1), CD3 (SK7), CD4 (RPA-T4), CD8 (RPA-T8), IFN-γ (4SB3), TNF-α (MAb11), were purchased from Biolegend (USA). For immunoblotting, the following antibodies, i.e., anti-IRF3 mAb (A19717, ABclonal, China), anti-Phospho-IRF3 mAb (4947, CST, USA), anti-TBK1 mAb (A3458, ABclonal), anti-Phospho-TBK1 mAb (5483, CST), and anti-β-actin mAb (AC004, ABclonal) were used. The antibodies used for immunofluorescence were anti-DNA mAb (CBL186, Sigma, USA), anti-TOM20 mAb (A19403, ABclonal), AF488-conjugated Goat anti-Rabbit (AS073, ABclonal), AF594-conjugated Goat anti-Mouse (AS054, ABclonal). Additional reagents included monomethyl fumarate (HY-103252, MCE, China) and dimethyl fumarate (HY-17363, MCE).

### Mouse model

For DMF treatment model, DMF was dissolved in dimethyl sulfoxide (DMSO) and solubilized under ultrasonic conditions at 37°C. The resulting solution was sequentially mixed with 10% DMSO, 40% PEG300, 5% Tween-80 and 45% saline, ensuring thorough homogenization prior to use for oral gavage administration. For in vivo experiments, 1 × 10⁵ TC-1 cells were subcutaneously implanted into C57BL/6J wild-type (WT) or BALB/c nude mice. DMF treatment (30 mg/kg) was initiated via oral gavage on day 7 post-inoculation and continued daily for 10 consecutive days. For anti-PD-1 combination therapy, anti-PD-1 antibodies (100 μg per mouse) were administered intraperitoneally on day 10, 13, and 16. For CD4/CD8 depleting model, TC-1 tumor-bearing mice were treated with 100 μg of CD4- or CD8-depleting antibodies on day 3, 6, 9, and 12 post-tumor inoculation.

For in vivo transplant tumor model with inoculation of DMF/MMF-treated TC-1, TC-1 cells were treated with 100 μM MMF or 50 μM DMF for 24 h and then subcutaneously implanted into C57BL/6J WT or BALB/c nude mice. For animal model examining the immunogenicity of DMF/MMF-treated TC-1 cells, mice that failed to develop tumors after subcutaneous implantation of DMF-pretreated TC-1 cells and control mice injected with PBS, were implanted with untreated TC-1 cells into the contralateral side 30 days later.

For PDX model, the patient-derived tumor tissues and paired TILs were established previously (25). Tumor tissue blocks with a volume of 3-4 mm³ were implanted into NCG mice. Treatment started on day 8 of tumor growth with intravenous adoptive transfer of TILs at a dose of 1 × 10⁷ cells per mouse, administered once every seven days, followed by intraperitoneal injection 45000 IU IL-2 at the same time as TILs, continuing for three consecutive days and then one-day intervals. On day 15 of tumor growth, oral gavage treatment with 30 mg/kg DMF was initiated, administered once daily for 10 consecutive days.

### RNA-seq

TC-1 cells were treated with 50 µM DMF or DMSO for 24 h. RNA was extracted from these samples and sent to BGI (China, Shenzhen) for sequencing. The cDNA library was constructed using the BGISEQ-500 platform. After obtaining clean reads, they were aligned to the reference genome sequence (GCF_000001635.26_GRCm38.p6) using HISATs. Differential gene expression analysis, Gene ontology (GO) enrichment analysis, Gene set enrichment analysis (GSEA) and heat map analysis were performed by the Dr. Tom online system (BGI-Shenzhen, China).

### qRT-PCR

Total RNA was extracted from cells with FastPure Cell/Tissue Total RNA isolation Kit (RC112, Vazyme, China) and reverse transcribed into cDNA by using HiScript II Reverse Transcriptase (R201, Vazyme). cDNA was used as a template and qPCR was performed using ChamQ Universal SYBR qPCR Master Mix kit (Q711, Vazyme) on a Bio-Rad CFX Connect Real-Time PCR System (v.2.0). The primer sequences were described in **Supplementary Table 1**.

### Quantification of mtDNA in cytoplasm

For the relative quantification of cytosolic mtDNA, the cytosolic fraction was isolated using the Mitochondria/Cytosol Fractionation Kit (K256-25, Biovision, China). DNA was extracted from whole-cell and cytosolic fractions using the Microsample Genomic DNA Extraction Kit (DP316, TIANGEN, China). qPCR was performed on whole-cell extracts and cytosolic fractions using mtDNA or nucDNA primers (**Supplementary Table 2**). The Ct values obtained from the DNA of whole-cell extracts were used as a normalization control for the DNA values obtained from cytosolic fractions, enabling effective sample standardization.

### Preparation of tumor single cell suspension

Mouse tumor tissues were minced and digested in 2 ml of DMEM containing 1 mg/ml Collagenase D (11088866001, Roche, Switzerland), 10 μg/ml DNase (abs47047435, Absin, China), and 1% FBS at 37°C with shaking at 80 rpm for 1 h. After filtration through a 70 μm cell strainer, red blood cells were lysed using red blood cell lysis buffer. The remaining single-cell suspension was used for flow cytometric analysis or magnetic-activated cell separation (MACS).

### Naive CD8^+^ T cells activation

Murine CD8^+^ T cells were isolated from spleens by positive selection using naive CD8α^+^ T cell Isolation Kit (130-096-543, Miltenyi, Germany). Naive CD8^+^ T cells were treated with 100 μM MMF or 50 μM DMF, and simultaneously cultured with Dynabeads (11452D, Gibco, USA) in Lymphocyte culture medium (**Supplementary Table 3**) for 72 h.

### Flow cytometric analysis

For flow cytometry staining of single-cell suspensions from tumor tissues, cells were first stained with a Fixable Viability Staining dye (564406, BD, USA) at 4 °C for 20 min to identify dead cells and with Fc blocker for 15 min to prevent non-specific staining. To analyze surface markers, cells were incubated for 30 min at 4°C with a mixture of fluorescein-labeled antibodies. For intracellular cytokine staining, cells were stimulated with Cell Activation Cocktail (with Brefeldin A) (423304, Biolegend) for 5 h at 37°C. The cells were collected, fixed and permeabilized after staining of cell surface markers, then stained with the IFN-γ and TNF-α. Flow cytometry was carried out with the FACSVerse (BD).

### Transwell assay

TC-1 cells with vector control or *Ccl5*&*Cxcl10* KO were treated with 100 μM MMF or 50 μM DMF for 24 h, followed by drug withdrawal. The treated tumor cells were harvested, and equal numbers of cells were reseeded. Supernatants were collected over the following 24 h and placed in the lower chamber of a Transwell system (3421, Corning, USA), with activated CD8⁺ T cells added to the upper chamber. The number of migrated CD8⁺ T cells in the lower chamber was counted after 6 h.

### Immunoblotting

Cells were lysed in RIPA buffer containing protease and phosphatase inhibitors on ice for 30 min. The lysate was centrifuged at 10,000 × g for 20 min, and the supernatant was collected. Protein concentration was quantified using a BCA Protein Assay Kit (P0012, Beyotime, China), and samples were normalized to equal concentrations with SDS loading buffer. Proteins were denatured by boiling at 100°C for 4 min, separated via 10% SDS-polyacrylamide gel electrophoresis (SDS-PAGE), and transferred to Immun-Blot PVDF membranes (1620177, Bio-Rad, USA). Membranes were blocked with 5% bovine serum albumin (BSA) in TBST (Tris-buffered saline with 0.1% Tween-20) at room temperature for 1 h, incubated with primary antibodies at 4°C overnight, and washed five times with TBST (5 min per wash). Subsequently, membranes were incubated with secondary antibodies at room temperature for 2 h, washed five times with TBST, and protein bands were visualized using ECL chemiluminescent reagent (HY-K1005, Mlbio, China).

For quantification of phosphorylated proteins (e.g., p-TBK1, p-IRF3), after detecting phosphorylated proteins, membranes were stripped with Stripping Buffer (CW0056M, Cwbio,China) at room temperature for 15 min to remove bound antibodies, re-blocked, and incubated with antibodies against total proteins (e.g., TBK1, IRF3), followed by repeated ECL detection. Band intensities were analyzed by ImageJ software (v.1.52), and phosphorylated protein expression levels were normalized to the corresponding total proteins.

### Measurement of cytokine levels

For cell culture supernatant, tumor cells were treated with 100 μM MMF or 50 μM DMF for 24 h, followed by drug withdrawal. The treated tumor cells were harvested, and equal numbers of cells were reseeded. Supernatants were collected after another 24 h of incubation. For tumor interstitial fluid, the tumor tissue is placed on a 70 µm filter secured at the top of a 15 ml centrifuge tube. The sample was centrifuged at 4°C, 2000 × *g* for 20 min, and the filtrate collected from the lower layer was collected. The levels of CXCL10 and CCL5 from supernatant or tumor interstitial fluid were measured using the mouse ELISA kit (88-56009, Invitrogen, USA and DY466-05, R&D, USA) according to the manufacturer’s instructions.

### Measurement of cGAMP

To measure 2’3’-cGAMP content, cells were treated with 100 μM MMF or 50 μM DMF for 24 h, and were then collected and washed with cold PBS, followed by adding 50 µl of PBS for every 1 × 10^5^ cells. Subsequently, cells were subject to three cycles of freeze-thaw procedure by freezing at -80°C for 30 mins and then thawing rapidly in 37°C water bath, to disrupt cell membranes. Cell debris was removed and the supernatant was collected as the cell lysate after centrifuge the thawed cells at 12,00 × for 10 mins at 4°C. The levels of 2’3’-cGAMP from cell lysates were measured using the 2’3’- cGAMP ELISA Kit (501700, Cayman, USA) according to the manufacturer’s instructions. The control was set to 1.

### Immunofluorescence microscopy

Cells grown on glass slides were fixed with 4% paraformaldehyde, then permeabilized with 0.1% Triton X-100. After blocking with 5% BSA for 1 h at room temperature and incubating with primary antibodies overnight at 4 °C, cells were washed with PBS and incubated with fluorophore-conjugated secondary antibodies for 1 h at room temperature and then washed with PBS. Super-resolution imaging was performed using commercialized High Intelligent and Sensitive SIM (HIS-SIM) provided by Guangzhou CSR Biotech Co. Ltd. Images were acquired using a 100 × /1.5 NA oil immersion objective.

### mtDNA depletion

To induce mtDNA depletion in tumor cells, the cells were cultured in DMEM containing 10% FBS, with the addition of 100 ng/mL ethidium bromide (HY-D0021, MCE) or 100 nM 2’,3’-dideoxycytidine (HY-17392, MCE). Culture medium was replaced every 2 days for 20 days.

### Culture of PDXO

The establishment of PDX-derived organoid (PDXO) has been described in detail in previous studies (25). Briefly, PDXO was cultured at 37°C in PDXO culture medium (**Supplementary Table 4**). Under these culture conditions, the transcription level of indicated molecules in PDXOs treated with MMF/ DMF for 24 h was measured by qRT-PCR.

### Statistical analysis

Data are presented as mean ± SD. Statistical analyses for in vivo and in vitro assays involving multiple-group comparisons were conducted using Dunnett’s multiple comparisons test (one-way ANOVA), Sidak’s multiple comparisons test (two-way ANOVA), or Tukey’s multiple comparisons test (two-way ANOVA). For single comparisons, unpaired or paired two-tailed Student’s t-tests were applied as appropriate. In cases of non-Gaussian distribution, nonparametric tests were employed, including the Wilcoxon matched-pairs signed-rank test for paired single comparisons, the Mann– Whitney U test for unpaired single comparisons, and the Kruskal–Wallis test for unpaired multiple-group comparisons. All statistical analyses were performed using GraphPad Prism software (v.8). A *P* value of < 0.05 was considered statistically significant throughout the study.

## Results

### 1. DMF enhances anti-CC immunity through CD8^+^ T Cell-mediated mechanisms

Previous studies have shown that DMF treatment can result in tumor regression and/or reduced metastasis in several types of cancers (26). In the quest for validating these findings in CC and exploring the possible mechanisms, we first examined the therapeutic effect of DMF on transplant tumor in immunodeficient (nude) mice generated by TC-1 cell line inoculation (Figure 1A). Of note, DMF was applied at a clinically equivalent dose (i.e., 30 mg per kg body weight) frequently used in various autoimmune mice models (27, 28). We did not note a significant change in tumor growth after DMF treatment in the immunodeficient mice (Figure 1B and C), although DMF displayed certain tumoricidal effect on TC-1 cells in vitro (Figure S1A). Interestingly, when the same experimental system applied to immunocompetent mice (Figure 1D), DMF treatment significantly inhibited tumor growth (Figure 1, E and F). These findings thus suggest that although DMF does have anti-CC potential, this effect is dependent on a fully functional immune system.

**Fig 1.**
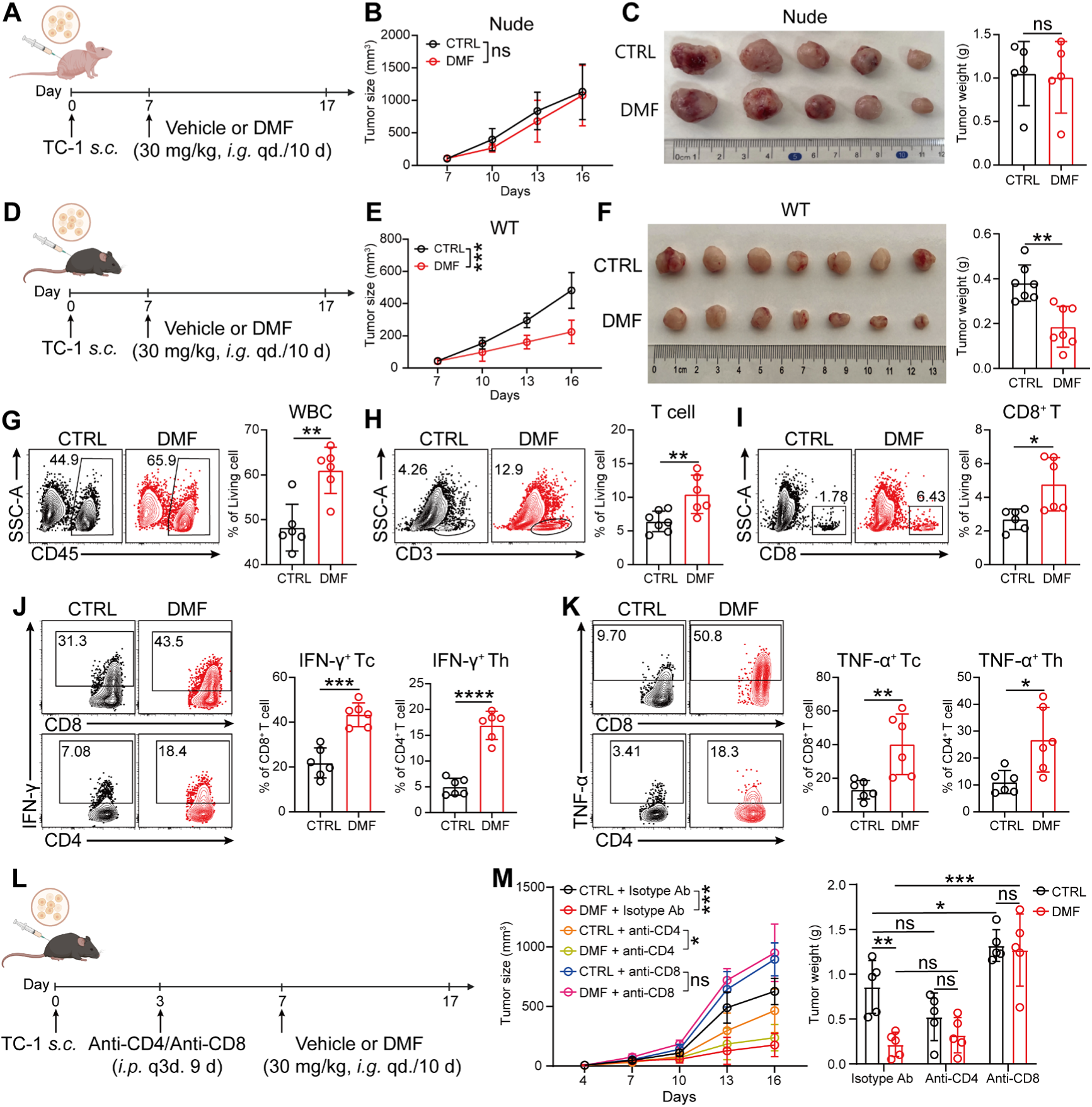
DMF enhances anti-CC immunity through CD8^+^ T Cell-mediated mechanisms. **A,** Schematic of experimental design for **B** and **C**. *i.g*., intragastric administration; *s.c*., subcutaneous. Created with BioRender.com. **B**, Tumor growth curve in TC-1 tumor-bearing nude mice following *i.g*. treatment with DMF. (n = 5 mice). **C**, Image of tumors (left) and weight measurements (right) in nude mice treated with or without 30 mg per kg body weight of DMF at day 17. **D,** Schematic of experimental design for **E** and **F**. Created with BioRender.com. **E**, Tumor growth curve in TC-1 tumor-bearing immunocompetent mice following *i.g*. treatment with DMF. (n = 7 mice). **F**, Image of tumors (left) and weight measurements (right) in immunocompetent mice treated with or without 30 mg per kg body weight of DMF at day 17. **G-K,** As in **D**, percentage of infiltrating white blood cells (WBC) (**G**); total T cells (**H**); CD8⁺ T cells (**I**); the proportions of IFN-γ⁺ (**J**) and TNF-α⁺ (**K**) cells among CD8⁺ T cells and CD4⁺ T cells in tumor of immunocompetent mice were analyzed by flow cytometry. (n = 6 mice). **L,** TC-1 tumor-bearing mouse treated with or without DMF and intraperitoneal (*i.p.*) injected with 100 μg of CD4/CD8-depleting antibodies on day 3, 6, 9, and 12 post tumor inoculation. Created with BioRender.com. **M,** Tumor growth curves (left) and tumor weights (right) were measured. (n = 5 mice). Data are the mean ± SD. *P* values were calculated using unpaired two-tailed Student’s *t* test (**B, C, E**-**K**), two-way ANOVA for Tukey’s multiple comparisons test (**M**). \**P* < 0.05, \*\**P* < 0.01, \*\*\**P* < 0.001, \*\*\*\**P* < 0.0001, and ns indicating no significant difference.

Immunophenotyping of tumor tissues from immunocompetent mice were performed using flow cytometry (Figure S1B). Compared to the control group, DMF treatment resulted in a significant increase in the number of infiltrating leukocytes in the tumor tissue (Figure 1G), with a particularly prominent increase in T cell infiltration (Figure 1H). Further analysis revealed a significant increase in CD8^+^ T cell infiltration (Figure 1I), whereas CD4^+^ T cells showed only minor changes (Figure S1C). Moreover, CD8^+^ T cell subsets with tumoricidal capacity, such as IFN-γ^+^CD8^+^ T cells (Figure 1J) and TNF-α^+^CD8^+^ T cells (Figure 1K), were also significantly elevated, suggesting that DMF may enhance immune-mediated anti-CC responses by recruiting/activating CD8^+^ T cells.

Next, to confirm the role of T cell subsets in the DMF-mediated antitumor effect, we selectively depleted CD8^+^ T cells and CD4^+^ T cells in the DMF treatment mouse models (Figure 1L and Figure S1D). The results showed that depletion of CD8^+^ T cells (Figure 1M) but not CD4^+^ T cells, significantly reversed the anti-CC effect of DMF, further highlighting the crucial role of CD8^+^ T cells in mediating DMF-induced anti-CC immune responses.

### 2. DMF enhances CD8^+^ T cell-mediated anti-CC immunity by improving the immunogenicity of tumor cells

We next sought to determine whether DMF acts directly on CD8^+^ T cells to exert anti-CC effects by employing in vitro experiment. DMF/MMF treatment significantly inhibited the proliferation of CD8⁺ T cells (Figure S2A). Moreover, although the cytotoxic proportion of CD8⁺ T cells increased after DMF/MMF treatment, their total number decreased significantly, resulting in a reduction in the absolute number of cytotoxic T cells (Figure S2, B and C). Therefore, the improved infiltration and anti-tumor function of CD8^+^ T cells caused by DMF treatment may result from an indirect mechanism.

Previous studies have shown that altering the immunogenicity of tumor cells can influence the function of immune cells within the tumor microenvironment (29, 30). Therefore, we asked if the antitumor function of DMF is exerted through similar mechanism. To test this, TC-1 cells were treated with DMF or MMF in vitro for 24 h and were subsequently inoculated into either immunodeficient or immunocompetent mice (Figure 2, A and D). Similar to the in vivo treatment model, DMF/MMF-treated and untreated TC-1 cells showed no significant growth difference in immunodeficient mice. (Figure 2, B and C). However, in immunocompetent mice, tumor growth was markedly inhibited, with some mice in the DMF -pretreated group failing to develop tumors (Figure 2, E and F). Furthermore, the tumors that were significantly suppressed in growth were accompanied by increased infiltration of CD8⁺ T cells with enhanced effector function (Figure 2, G-L). These results suggest that DMF indirectly promotes CD8^+^ T cell-mediated anti-tumor immunity through acting on tumor cells.

**Fig 2.**
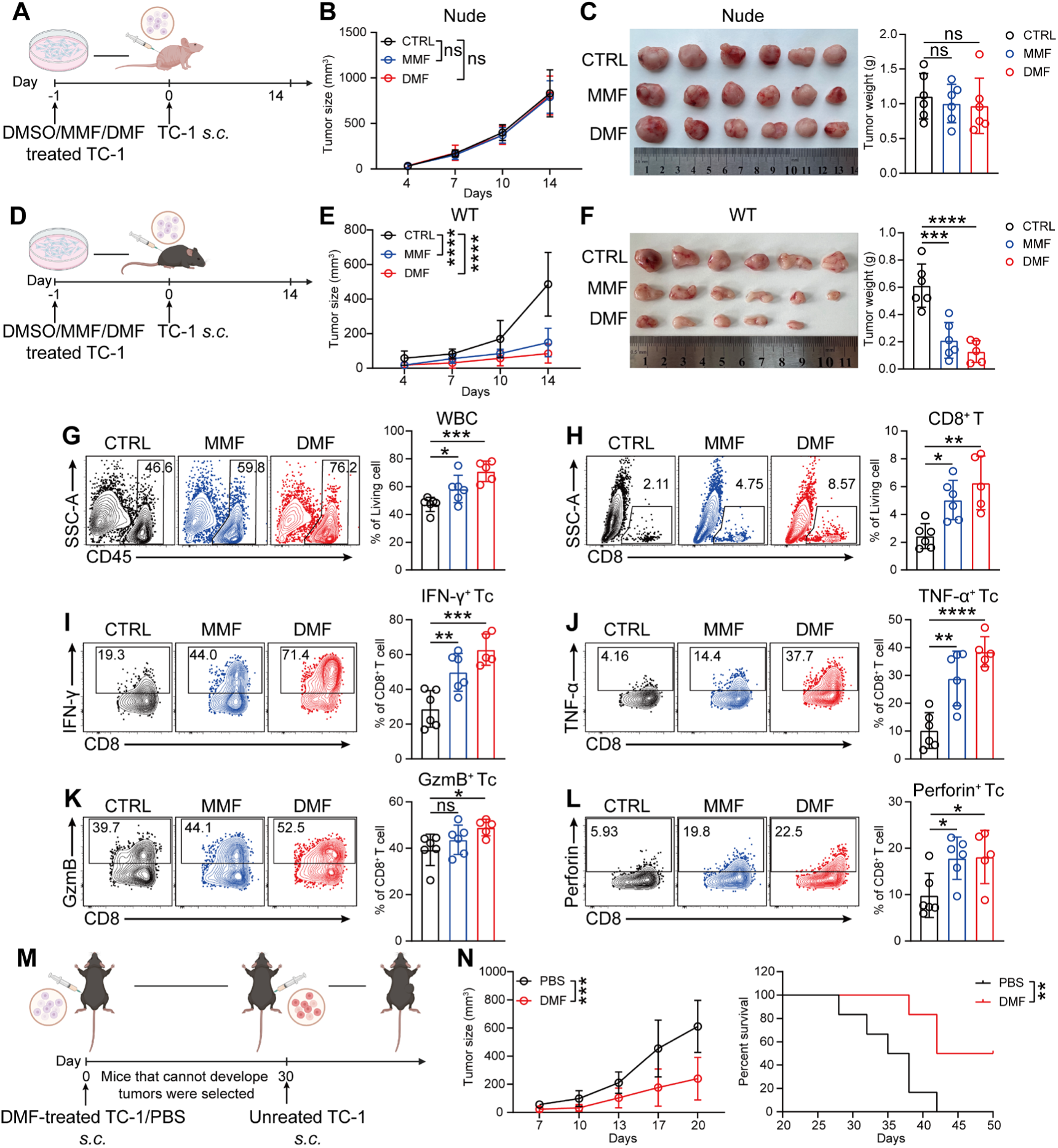
DMF enhances CD8^+^ T cell-mediated anti-CC immunity by improving the immunogenicity of tumor cells. **A,** Schematic of experimental design for **B** and **C**. Created with BioRender.com. **B**, Tumor growth curves. **C**, Image of tumors (left) and weight measurements (right) in immunodeficient mice. (n = 6 mice). **D,** Schematic of experimental design for **E** and **F**. Created with BioRender.com. **E**, Tumor growth curves. **F**, Image of tumors (left) and weight measurements (right) in immunocompetent mice. (n = 6 mice). **G-L**, As in **D**, percentage of WBC (**G**); CD8⁺ T cells (**H**); IFN-γ⁺CD8⁺ T cells (**I**); TNF-α⁺CD8⁺ T cells (**J**); GzmB⁺CD8⁺ T cells (**K**) and Perforin⁺CD8⁺ T cells (**L**) in immunocompetent mice were analyzed by flow cytometry. (n = 6 mice). **M**, Schematic of experimental design for **N**. Created with BioRender.com. **N**, Tumor growth curves (left) and mouse survival curves (right) were recorded. (n = 7 mice). Data are the mean ± SD. *P* values were calculated using one-way ANOVA for Dunnett’s multiple comparisons test (**C**, **F**, **G**-**L**), two-way ANOVA for Tukey’s multiple comparisons test (**B, E, N** left) and log-rank test for survival analysis (**N** right). \**P* < 0.05, \*\**P* < 0.01, \*\*\**P* < 0.001, \*\*\*\**P* < 0.0001, and ns indicating no significant difference.

Next, we interrogate whether DMF treatment enhances the immunogenicity of tumor cells by utilizing a tumor-reinoculation mouse model. For this, mice from the DMF-pretreated group that did not develop tumors were reinoculated with tumor cells into the contralateral flank 30 days post-initial inoculation (Figure 2M). The results showed that, compared to the group injected with PBS on the left side, mice with DMF-pretreated TC-1 cells that failed to form tumors on the left side exhibited significantly slower tumor growth on the right side and had markedly higher survival rates (Figure 2N). Collectively, these findings suggest that pretreatment of tumor cells with DMF induces a potent anti-tumor immune response through improving the immunogenicity of tumor cells.

### 3. DMF enhances CD8^+^ T cell infiltration through inducing tumor cells to secrete CCL5 and CXCL10

To further explore the underlying mechanism, we performed RNA-seq on DMF-treated and -untreated TC-1 cells for differential gene expression analysis. The volcano plot of differentially expressed genes (DEGs) showed that *Cxcl10*, *Ifit3*, *Ifit1*, and *Ccl5*, which are associated with the Type I interferon pathway (31, 32), were significantly upregulated in DMF-treated tumor cells (Figure 3A). Consistently, gene set enrichment analysis (GSEA) also indicated an upregulation of the Type I interferon pathway (Figure 3B). Moreover, a comparative analysis of DEGs between the DMF-treated and -untreated groups showed a marked upregulation of multiple type I interferon-responsive genes following DMF treatment (Figure 3C), which were further confirmed by qRT-PCR (Figure 3D). Additionally, when the human CC cell line SiHa was selected to validate this finding, DMF/MMF treatment again led to upregulation of the *CCL5* and *CXCL10* genes (Figure 3E). Moreover, higher levels of CCL5 and CXCL10 were detected in the supernatant of DMF-treated TC-1 cells, further supporting these findings (Figure 3F). Importantly, tumor cells extracted from DMF-treated mice (Figure S2D) exhibited significantly upregulated expression levels of *Ccl5* and *Cxcl10* compared to those from untreated mice, as determined by qRT-PCR analysis (Figure 3G). Measurement of CCL5 and CXCL10 levels in the tumor interstitial fluid revealed significantly higher concentrations of both chemokines in the DMF-treated group (Figure 3H). Therefore, our in vitro and in vivo findings together suggest that the heightened type I interferon response, especially increased release of CCL5 and CXCL10 by tumor cells may be responsible for DMF treatment-induced CD8^+^ T cell infiltration.

**Fig 3.**
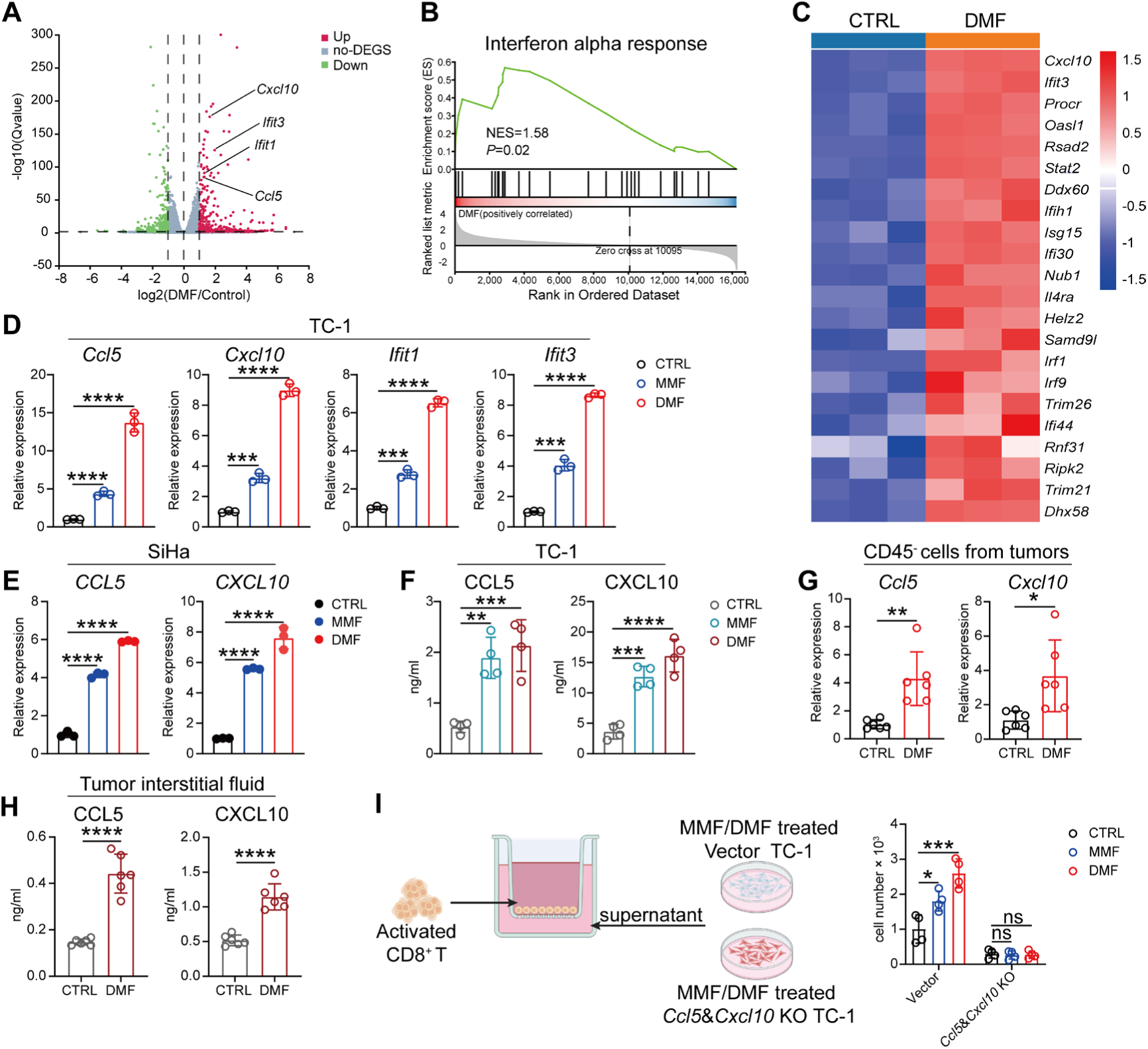
DMF enhances CD8^+^ T cell infiltration through inducing tumor cells to secrete CCL5 and CXCL10. **A-C**, RNA-seq results of TC-1 cells treated with or without 50 µM DMF for 24 h. **A**, Volcano plot of differentially expressed genes (DEGs), type I interferon response relative genes were annotated. **B**, GSEA plot showing the interferon alpha response pathway. **C,** Heatmap of DEGs related to the type I interferon response. **D**, Transcriptional levels of *Cxcl10*, *Ccl5*, *Ifit1*, and *Ifit3* in TC-1 cells treated with or without 100 μM MMF or 50 μM DMF for 24 h, analyzed by qRT-PCR. **E**, Transcriptional levels of *CXCL10* and *CCL5* in SiHa cells treated with or without 100 μM MMF or 50 μM DMF for 24 h, analyzed by qRT-PCR. **F**, Concentrations of CXCL10 and CCL5 in the supernatant of TC-1 cells after 24 h of treatment with 100 μM MMF or 50 μM DMF, quantified by ELISA. **G** and **H**, After 10 days of treatment with DMF by gavage, tumors were harvested from immunocompetent mice. (as in Fig. 1D). **G**, Transcriptional levels of *Cxcl10* and *Ccl5* in CD45⁻ cells derived from tumor tissue, analyzed by qRT-PCR. (n = 6 mice). **H**, Concentrations of CXCL10 and CCL5 in the tumor interstitial fluid, quantified by ELISA. (n = 6 mice). **I**, Schematic of experimental design for the Transwell system (left) created with BioRender.com and the number of migrated CD8⁺ T cells in the lower chamber was counted after 6 h (right). Data are the mean ± SD. *P* values were calculated using unpaired two-tailed Student’s *t* test (**G** and **H**), one-way ANOVA for Dunnett’s multiple comparisons test (**D**-**F**), two-way ANOVA for Tukey’s multiple comparisons test (**I right**). \**P* < 0.05, \*\**P* < 0.01, \*\*\**P* < 0.001, \*\*\*\**P* < 0.0001, and ns indicating no significant difference.

CCL5 and CXCL10 are well-known for their capacity to recruit CD8^+^ T cells into tissues including tumors (30, 33). Indeed, CCL5 and CXCL10 were the top two chemokines with increased expression in tumor cells after DMF treatment (Figure S2E). Thus, we went on to determine whether these two chemokines are required for DMF-induced CD8^+^ T cell infiltration. To this end, we first created *Ccl5-* and *Cxcl10*-knockout (KO) TC-1 cell line and confirmed the lack of these chemokines in the supernatant, respectively (Figure S2F). Then, supernatant from knockout and vector cell lines were collected after DMF/MMF-treatment and placed in the lower chamber of a transwell system, followed by examining their capacity to recruit activated CD8^+^ T cells in the upper chamber after 6 h of incubation. The results showed that although supernatants from vector tumor cell significantly recruited more CD8^+^ T cells into the lower chamber when tumor cells were treated with DMF/MMF (Figure 3I), those from *Ccl5* and *Cxcl10* KO cell line failed to do so, indicating that CCL5 and CXCL10 are required for DMF/MMF-treated CC cells to promote CD8^+^ T cell recruitment.

### 4. cGAS-STING-TBK1 pathway mediates the DMF-induced anti-tumor immune response

cGAS-STING-TBK1 pathway has been reported to be pivotal in inducing type I interferon response in the fight against infectious agents and tumor (34). Therefore, we next explored whether the DMF/MMF-induced anti-CC immune response is regulated by this pathway. Indeed, TC-1 cells exhibited significantly increased expression of p-TBK1 and p-IRF3 (Figure 4A), along with elevated intracellular cGAMP levels (Figure 4B) after DMF/MMF treatment. Next, we repeated the experiment in the human-derived SiHa CC cell line and obtained similar results (Figure 4, C and D), indicating that the capacity of DMF/MMF to activate type I interferon response is conserved across CC cells from different sources.

**Fig 4.**
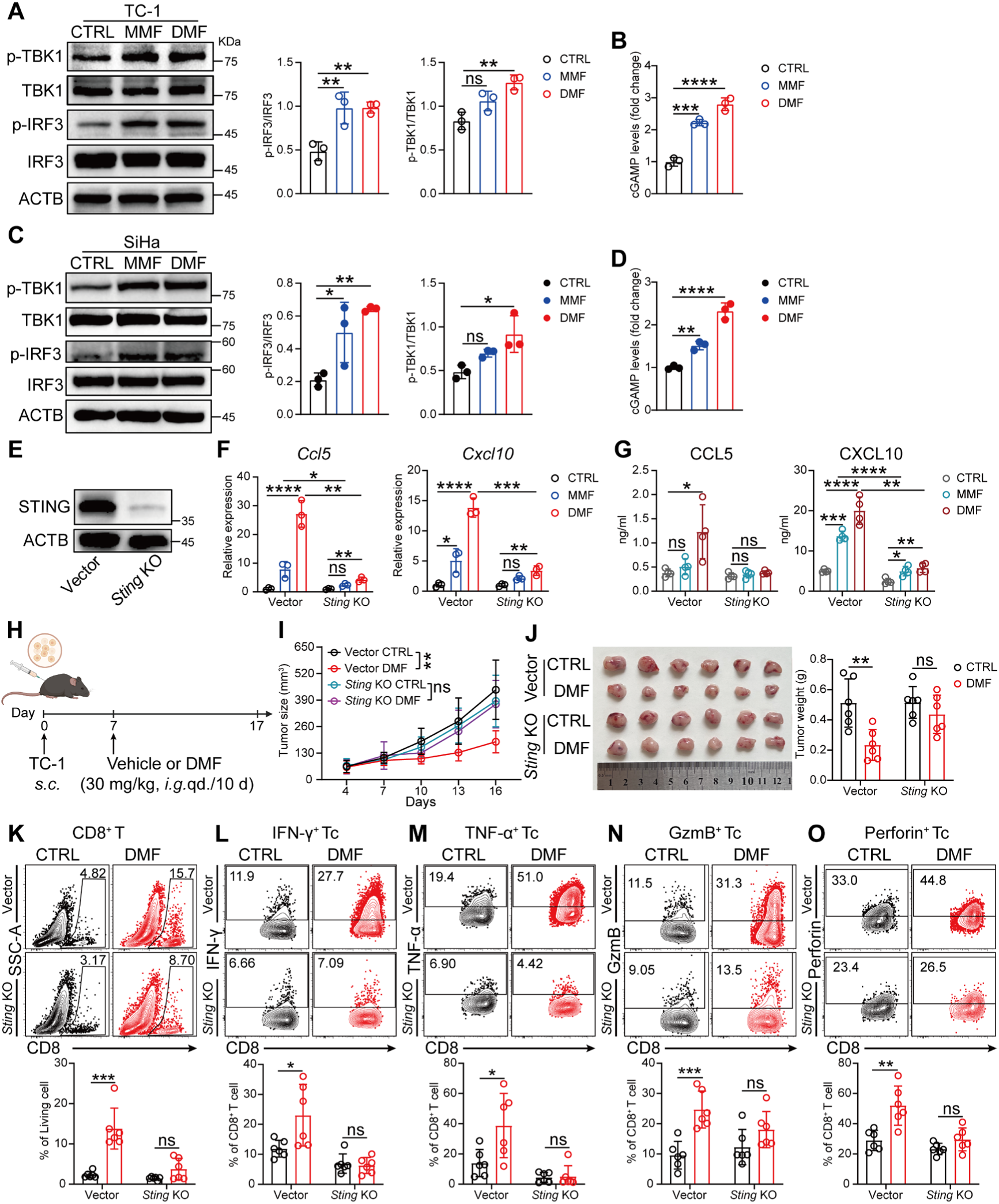
cGAS-STING-TBK1 pathway mediates the DMF-induced anti-tumor immune response. **A** and **B**, Immunoblots of IRF3, p-IRF3, TBK1, p-TBK1 (**A**), and relative abundance of cGAMP (**B**) in TC-1 cells treated with 100 μM MMF or 50 μM DMF for 24 h. **C** and **D**, Immunoblots of IRF3, p-IRF3, TBK1, p-TBK1(**C**), and relative abundance of cGAMP (**D**) in SiHa cells treated with 100 μM MMF or 50 μM DMF for 24 h. **E**, Immunoblots of STING in TC-1 cells, comparing Vector and *Sting* knockout (KO) cells. **F** and **G**, CCL5 and CXCL10 were quantified by qRT-PCR (**F**) and ELISA (**G**) respectively, in vector or *Sting* KO TC-1 cells treated with 100 μM MMF or 50 μM DMF for 24 h. **H**-**O**, As in Fig. 1A, a total of 1 × 10⁵ vector or *Sting*^-/-^ TC-1 cells were subcutaneously implanted into mice. **H**, Schematic of experimental design created with BioRender.com. **I**, Tumor growth curves. **J**, Image of tumors (left) and weight measurements (right) at day 17. **K-O**, Percentage of infiltrating CD8⁺ T cell populations, including total CD8⁺ T cells (**K**), IFN-γ⁺CD8⁺ T cells (**L**), TNF-α⁺CD8⁺ T cells (**M**), GzmB⁺CD8⁺ T cells (**N**), and Perforin⁺CD8⁺ T cells (**O**) in tumor were analyzed by flow cytometry. (n = 6 mice). Data are presented as the mean ± SD. *P* values were calculated using one-way ANOVA for Dunnett’s multiple comparisons test (**A**-**D**), two-way ANOVA for Tukey’s multiple comparisons test (**F** and **G, I-O**). \**P* < 0.05, \*\**P* < 0.01, \*\*\**P* < 0.001, \*\*\*\**P* < 0.0001, and ns indicating no significant difference.

Since type I interferon response can be induced through pathways independent of STING (e.g., TLRs), we therefore tested if STING is required for DMF/MMF to activate type I interferon response by generating a *Sting* KO TC-1 cell line (Figure 4E). The results showed that in *Sting*-deficient TC-1 cells, the upregulation of type I interferon-related genes after DMF/MMF-treatment was significantly reduced compared to that in vector cells (Figure 4F, Figure S3A). Similarly, the levels of CCL5 and CXCL10 in the culture supernatants were also decreased (Figure 4G). Subsequently, we implanted *Sting* KO TC-1 cells into mice and treated the tumor-bearing mice with DMF (Figure 4H). In contrast to that observed in WT group, DMF treatment did not significantly affect tumor size in *Sting* KO group (Figure 4, I and J). Moreover, the proportion and function of CD8^+^ T cells infiltrating the tumor tissues remained unchanged (Figure 4, K-O). Together, these results indicate that the cGAS-STING-TBK1 pathway plays an indispensable role in DMF-induced antitumor immune responses.

### 5. DMF/MMF treatment induces tumor cells to release mtDNA into the cytoplasm

We went on to investigate the potential mechanisms underlying the upregulation of the cGAS-STING pathway in tumor cells following DMF/MMF treatment. GSEA analysis revealed that genes related to oxidative stress response were significantly enriched in DMF-treated TC-1 cells compared to the control group (Figure 5A). Further experiments showed that DMF/MMF treatment significantly increased intracellular ROS levels (Figure 5B) and decreased mitochondrial membrane potential (Figure 5C), indicating potential mitochondrial damage. Electron microscopy analysis further confirmed mitochondrial cristae disorganization in tumor cells following DMF/MMF treatment (Figure 5D).

**Fig 5.**
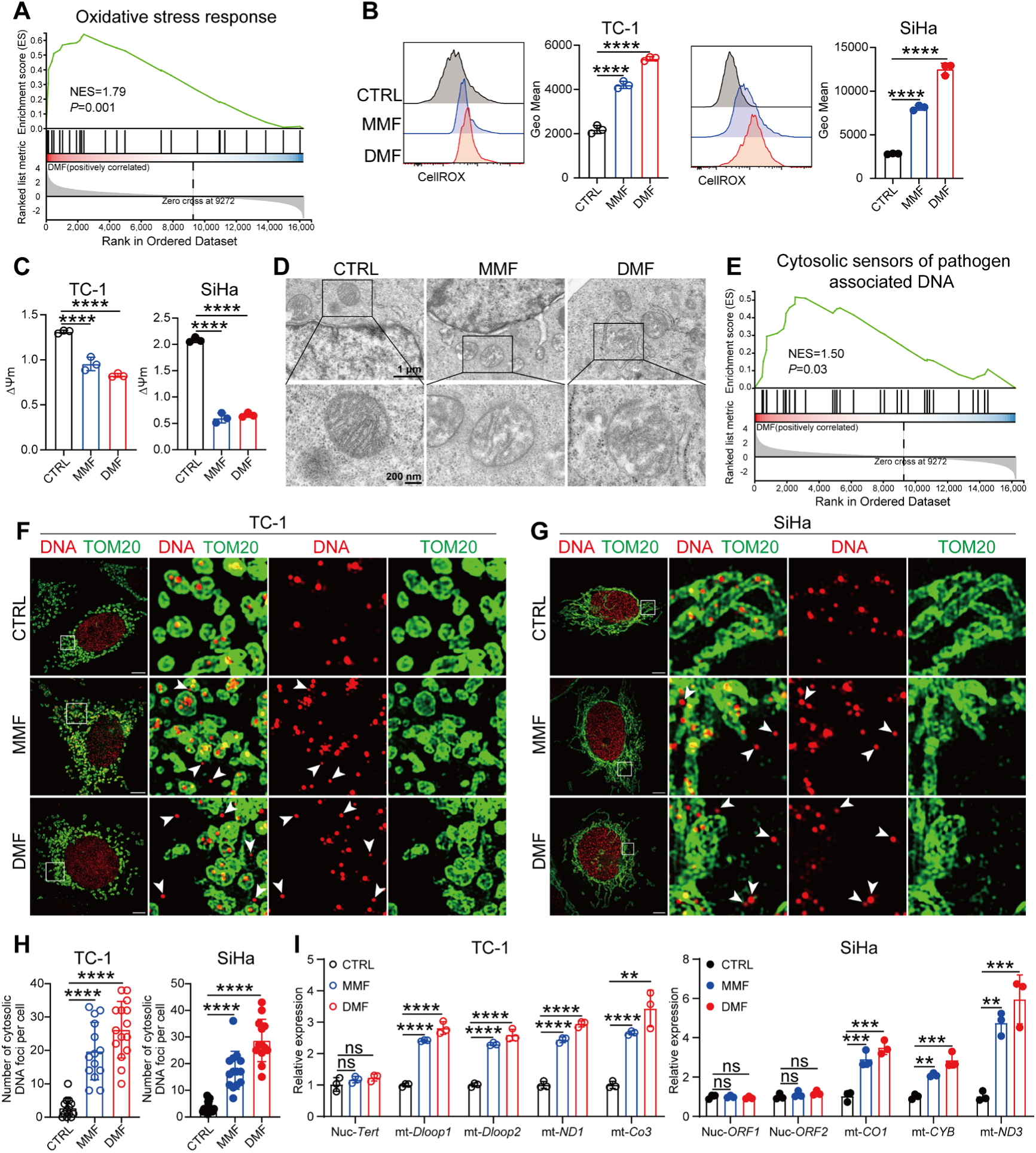
DMF/MMF treatment induces tumor cells to release mtDNA into the cytoplasm. **A**, GSEA plot of the oxidative stress response in TC-1 cells treated with or without 50 μM DMF. **B** and **C**, CellROX levels (**B**) and mitochondrial membrane potential (**C**) were measured in TC-1 and SiHa cells treated with 100 μM MMF or 50 μM DMF for 12 h. **D**, Electron microscopy of TC-1 cells treated with 100 μM MMF or 50 μM DMF for 24 h. **E**, GSEA plot of the Cytosolic sensors of pathogen associated DNA pathway in TC-1 cells treated with or without 50 μM DMF. **F**-**H**, Immunofluorescence image of tumor cells treated with 100 μM MMF or 50 μM DMF for 24 h, (**F**) showing TC-1 cells, (**G**) showing SiHa cells. DNA, red; TOM20, green; scale bar = 5 μm. **H**, Quantitative analysis of extramitochondrial DNA in the cytosol in figure **F** and **G**. (n = 13-15 cells). **I**, Relative expressions of nuclear (nuc) and mitochondrial (mt) DNA outside the mitochondria in the cytoplasm was detected by qRT-PCR. Data are presented as the mean ± SD. *P* values were calculated using one-way ANOVA for Dunnett’s multiple comparisons test (**B, C, H**), two-way ANOVA for Tukey’s multiple comparisons test (**I**). \**P* < 0.05, \*\**P* < 0.01, \*\*\**P* < 0.001, \*\*\*\**P* < 0.0001, and ns indicating no significant difference.

Additionally, GO enrichment analysis indicated a significant increase in cellular response to DNA damage stimulus pathway (Figure S3B), as well as a notable enrichment in the cytosolic sensors of pathogen-associated DNA pathway in the DMF-treated cells (Figure 5E). Based on these observations, we hypothesize that DMF/MMF-treatment may activate the cGAS-STING pathway by increasing the cytoplasmic DNA content. In line with this hypothesis, the cytoplasmic DNA content was significantly elevated after DMF/MMF treatment in both TC-1and SiHa cell lines, as shown by immunofluorescence staining (Figure 5F-H). To further verify the source of cytoplasmic DNA (i.e., nuclear or mitochondria) after DMF/MMF treatment, we excluded mitochondria and extracted the DNA from the cytoplasm for qPCR analysis. The results showed that the cytoplasmic DNA in both TC-1 and SiHa cells treated with DMF/MMF is primarily mtDNA rather than nuclear DNA (nucDNA) (Figure 5I). These results collectively suggest that DMF/MMF treatment induces mitochondrial damage, leading to an increase in cytoplasmic mtDNA content.

### 6. Cytoplasmic mtDNA is required for DMF/MMF-induced activation of the cGAS-STING pathway and antitumor response

We next investigated whether the release of mitochondrial DNA into the cytoplasm contributes to the activation of cGAS-STING pathway and the subsequent type I interferon response in DMF/MMF-treated cells by using an established protocol to deplete mtDNA. This was achieved by culturing cells in either low concentrations of ethidium bromide (EthBr) (35) or at the presence of 2′-3′-dideoxycytidine (DDC) (36) (Figure S3C). As expected, depletion of mtDNA significantly reduced the upregulation of type I interferon response-related genes induced by DMF/MMF treatment (Figure 6A, Figures S3, D and E), and also decreased the secretion levels of CCL5 and CXCL10 (Figure 6B). Additionally, mtDNA depletion in both TC-1 and SiHa cells suppressed DMF/MMF-induced activation of the cGAS/STING pathway (Figure 6, C-F). Importantly, when mtDNA-depleted TC-1 cells (TC-1ρ0) were implanted subcutaneously into mice (Figure 6G), the inhibited growth of tumors pre-treated with DMF observed in the TC-1 group was no longer noted (Figure 6, H and I). These findings indicate that the release of mitochondrial DNA into the cytoplasm is required for DMF-induced activation of the cGAS-STING pathway and the subsequent anti-CC immune response.

**Fig 6.**
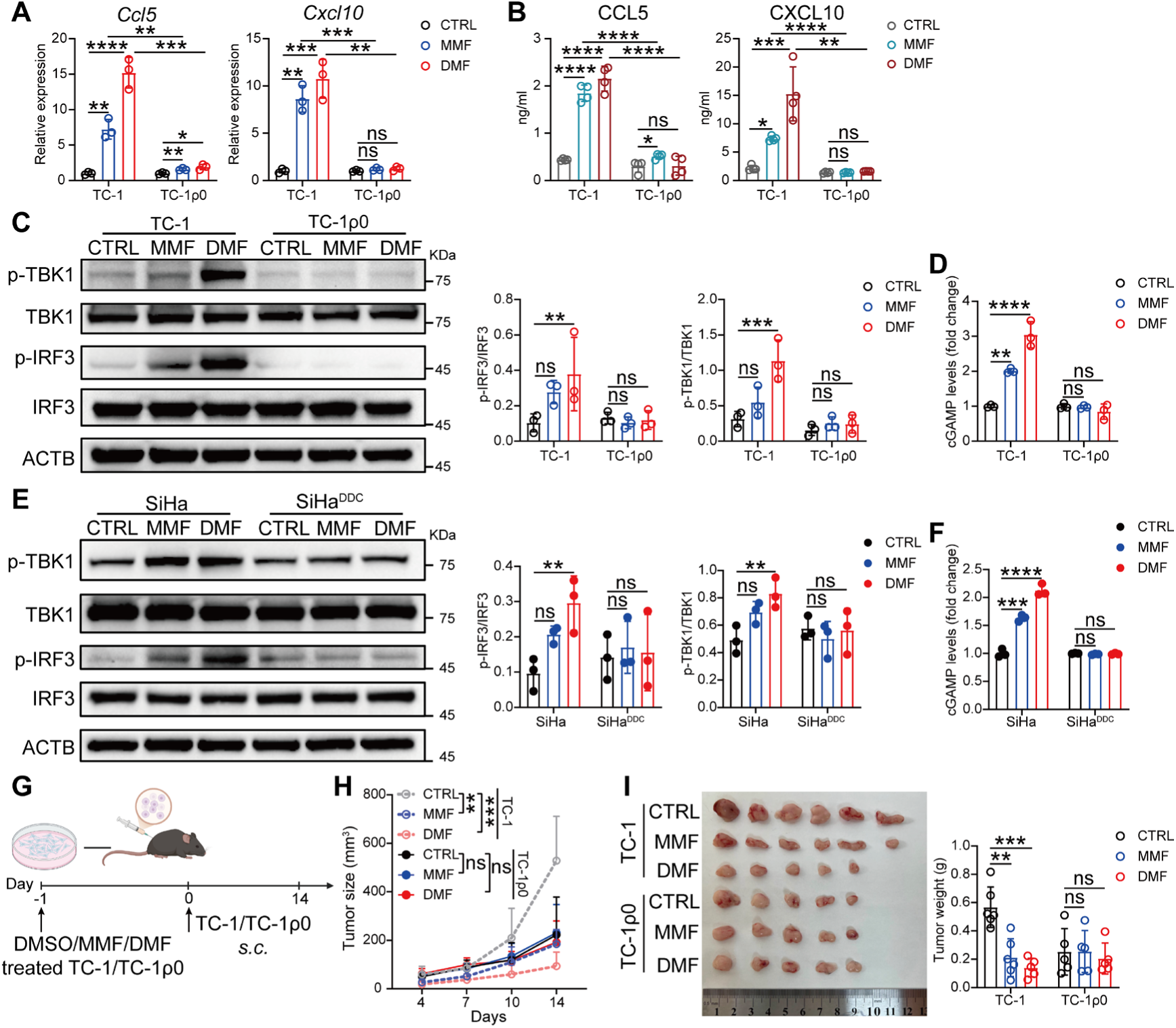
Cytoplasmic mtDNA is required for DMF/MMF-induced activation of the cGAS-STING pathway and antitumor response. **A** and **B**, CCL5 and CXCL10 were quantified by qRT-PCR (**A**) and ELISA (**B**) respectively, in TC-1 and TC-ρ0 cells treated with 100 μM MMF or 50 μM DMF for 24 h. **C** and **D**, Immunoblots of IRF3, p-IRF3, TBK1, p-TBK1 (**C**) and relative abundance of cGAMP levels (**D**) in TC-1 and TC-1ρ0 cells treated with 100 μM MMF or 50 μM DMF for 24 h. **E** and **F**, Immunoblots of IRF3, p-IRF3, TBK1, p-TBK1 (**E**) and relative abundance of cGAMP levels (**F**) in SiHa and SiHa^DDC^ cells treated with 100 μM MMF or 50 μM DMF for 24 h. **G**, Schematic of experimental design for **H** and **I**. Created with BioRender.com. **H**, Tumor growth curves. **I**, Image of tumors (left) and weight measurements (right) at day 15. (n = 5-6). Data are presented as the mean ± SD. *P* values were calculated using two-way ANOVA for Tukey’s multiple comparisons test (**A-F, H** and **I**). \**P* < 0.05, \*\**P* < 0.01, \*\*\**P* < 0.001, \*\*\*\**P* < 0.0001, and ns indicating no significant difference.

### 7. DMF improve the therapeutic efficacy of immunotherapy with PD-1 blockade and TILs

Although ICB therapy has been approved in CC, patients’ response to this therapy remains relatively low, which is partially attributable to the poor infiltration of ICB-reinvigorated CD8⁺ T cells (6, 37, 38). Given the capability of DMF treatment in enhancing the infiltration of CD8⁺ T cells, we therefore reasoned that DMF might be able to improve the efficacy of PD-1 blockade by promoting immune infiltration. Indeed, in the TC-1 transplant tumor model, monotherapy with anti-PD-1 antibody did not show significant efficacy, while DMF treatment alone already resulted in markedly prolonged mouse survival. Importantly, combinatorial therapy with DMF and anti-PD-1 antibody were superior to both monotherapies (Figure 7, A and B), indicating the synergistic effect of these two therapeutic regimens.

**Fig 7.**
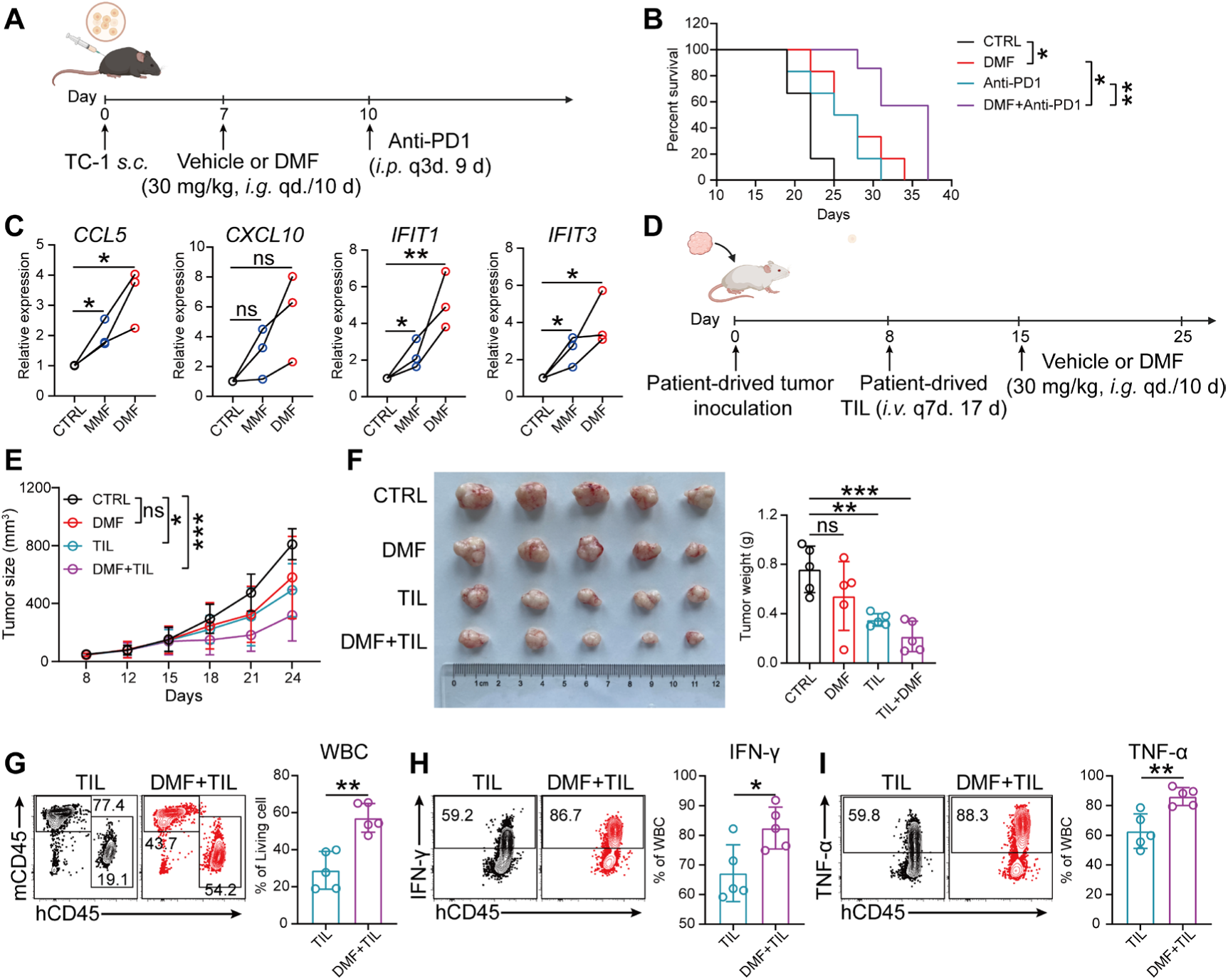
DMF improves the therapeutic efficacy of immunotherapy with PD-1 blockade and TILs. **A,** TC-1 tumor-bearing mouse gavaged with or without DMF and treated with 100μg of anti-PD1 antibodies on day 10, 13 and 16 post tumor inoculation. (n = 6 mice). Created with BioRender.com. **B**, Mouse survival curves were recorded. **C**, Transcriptional levels of *CXCL10*, *CCL*5, *IFIT1*, and *IFIT3* in PDXOs from cervical cancer patients treated with 100 μM MMF or 50 μM DMF, analyzed by qRT-PCR. (n = 3 patients). **D**, Schematic of experimental design for **E**-**F,** *i.v.* intravenous injections. *i.p.* intraperitoneal injections. Created with BioRender.com. **E**, Tumor growth curve. **F**. Image of tumors (left) and weight measurements (right) at day 25. The proportion of patient-derived WBCs (**G**), IFN-γ⁺ WBC cells (**H**) and TNF-α⁺ WBC cells (**I**) in PDX were analyzed. (n = 5 mice). Data are presented as the mean ± SD. *P* values were calculated using unpaired two-tailed Student’s *t* test (**G**-**I**), one-way ANOVA for Dunnett’s multiple comparisons test (**C**), two-way ANOVA for Tukey’s multiple comparisons test (**E** and **F**), log-rank test for survival analysis, and corrected by Bonferroni methods (**B**). \**P* < 0.05, \*\**P* < 0.01, \*\*\**P* < 0.001, \*\*\*\**P* < 0.0001, and ns indicating no significant difference.

We next asked if activating cGAS-STING pathway represents a feasible approach for treating human CC. To examine this issue, we first performed association analysis using the public gene database of CC patients, finding that high expression of *STING* is positively correlated with the prognosis of CC (Figure S3F), suggestive of activating cGAS-STING pathway as a promising therapeutic strategy in treating human CC. To validate this premise, we established PDX tumors from three CC patients and generated corresponding organoid samples (PDXO, Figure S3G) followed by testing the therapeutic efficacy of DMF toward these human-derived tumors. In line with that noted in murine TC-1 cells (Figure 3D), PDXO exhibited markedly increased expression of *CCL5*, *CXCL10*, *IFIT1* and *IFIT3* after DMF/MMF treatment (Figure 7C). Next, the in vivo evaluation of DMF/MMF’s capacity to induce anti-CC immune response against human tumors and hence tumor regression was performed in a PDX mouse model. As expected, daily treatment with DMF alone starting from day 15 after NCG mice were subcutaneously implanted with PDX tumor failed to inhibit PDX growth given the immunodeficiency status of NCG mice. Therefore, patient-derived tumor-infiltrating lymphocytes (TIL) from the same patient where PDX was derived were next prepared (Figure S3H) and infused at day 8, 15 and 22 post tumor inoculation into mice with or without DMF treatment (Figure 7D). The results showed that the combination of DMF and TIL effectively inhibited tumor growth, with the therapeutic effect being superior to that of TIL alone (Figure 7E, F). Flow cytometric analysis of tumor tissues revealed that DMF enhanced the infiltration and cytotoxic function of TILs (Figure 7G-I). Collectively, these findings suggest that combining DMF with PD-1 blockade or TIL therapy may offer a more effective treatment strategy by enhancing CD8⁺ T cell infiltration and function, thereby improving the clinical outcomes of human CC.

## Discussion

Developing effective immunotherapies is of paramount importance for treating CC, a common gynecological malignancy with high incidence and mortality rates (4). The findings of this study provide a novel solution by repurposing DMF, an FDA-approved drug previously used in treating autoimmune diseases, for the immunotherapy of CC. Through activating the cGAS-STING-TBK1 pathway in tumor cells, DMF enhances CD8⁺ T cell infiltration and modulates the tumor microenvironment, leading to CC regression. Considering the poor prognosis of patients with advanced CC and the limited efficacy of current immunotherapies, incorporating DMF into the existing treatment regimens may represent a promising strategy.

The direct anticancer activity of DMF has been documented dating back to 2006 in melanoma (39), and was later extended to breast cancer (23), colon cancer (40), ovarian cancer, and lung cancer (21). In this regard, multiple mechanisms have been suggested, including the inhibition of NF-κB signaling pathway, blocking tumor cell proliferation cycles, inducing apoptosis, and suppressing the expression of matrix metalloproteinase (18, 23, 24, 41). In CC, DMF has also been found to be capable of inhibiting the proliferation of human HeLa cells in vitro (41). In line with this, we did observe the direct tumor-suppressing activity of DMF toward murine CC cell line TC-1 in vitro. Although DMF at a low concentration (50 μM) demonstrated a modest in vitro tumor growth inhibition in TC-1 cells (Figure S1A), tumor growth was not affected in tumor-bearing immunodeficient mice after low-dose DMF treatment (Figure 1, A-C). Interestingly, however, the same DMF treatment regimen resulted in significant growth inhibition of implanted tumor in immunocompetent mice (Figure 1, E-F), suggesting that in addition to the direct anticancer activity, DMF can also indirectly result in tumor regression under the help of immune system, especially CD8^+^ T cells, as illustrated by the reverse of DMF-mediated tumor suppression upon CD8^+^ T cell-depletion (Figure 1, M). Therefore, the novel indirect anticancer activity of DMF uncovered by this study opens up a new horizon for the application of DMF in cancer therapy.

It is important to note that DMF was originally used as an anti-inflammation drug for treating patients with autoimmune diseases such as multiple sclerosis and psoriasis (14, 42), therefore might raise concerns about its systemic application in cancer patients, which could potentially compromise antitumor immune response. Indeed, DMF has been reported to inhibit autoreactive T helper (Th) 1 and Th17 cells in autoimmune disorders in both mouse models and human (43, 44). However, the effect of DMF on CD8^+^ T cells in the context of autoimmune diseases has been less-explored. A recent study by Lückel et al found in a murine experimental autoimmune encephalomyelitis (EAE) model that DMF treatment reduces the proportion of pro-inflammatory Tc17 cell subsets while driving the differentiation of CD8⁺ T cells toward a cytotoxic T lymphocyte (CTL) phenotype with enhanced cytotoxicity (45). In concordance with this observation, we found in the current study that naïve CD8^+^ T cells simultaneously exposed to activation stimuli (anti-CD3/CD28/IL-2) and DMF exhibited enhanced cytotoxicity but concurrently decreased proliferation compared to those exposed to activation stimuli alone (Figure S2, B-D). Although the differentiation status of CD8^+^ T cells used for DMF treatment in these two studies are different (Tc17 vs naïve), these results together suggest that CD8^+^ T cells may respond to DMF in a different way from Th cells, which only exhibit inhibited proliferation but not enhanced Th1 differentiation after DMF treatment. Indeed, the difference in metabolic requirements lies between CD8^+^ and CD4^+^ T cells (46), and DMF are known to be capable of regulating metabolism in immune cells (47, 48). However, further experiments are required to validate the above speculation.

In this study, both increased infiltration and enhanced cytotoxicity of CD8^+^ T cells in CC tissue were observed in tumor-bearing mice after DMF treatment (Figure 1, I-K). Although the direct cytotoxicity-promoting effect of DMF on CD8^+^ T cells cannot be excluded in this scenario as previously discussed, several lines of evidence suggest that these alterations are largely attributable to the indirect effect of DMF on CC cells. First, although in vitro treatment of CD8^+^ T cells with DMF did increase their cytotoxic differentiation, it also inhibited their proliferation, leading to the overall decreased CTL numbers, which is different from the observation in vivo. Second, when DMF-treated and -untreated TC-1 cells were implanted into immunocompetent mice, an experimental system in which CD8^+^ T cells were not exposed to DMF, similar alterations CD8^+^ T cells were observed in the DMF-treated but not -untreated group (Figure 2, G-L). Finally, elimination of mtDNA, STING, and CCL5/CXCL10 in DMF-treated TC-1 cells counteracted these effects. Therefore, these evidences, together with the well-known activity of cGAS-STING-IFN I pathway in promoting CTL differentiation, lend a strong support for the hypothesis that DMF promotes CD8^+^ T cell-mediated antitumor response and thereby tumor regression through activating mtDNA-cGAS-STING-IFN I-CCL5/CXCL10 pathway in tumor cells.

Although the pivotal role of cGAS-STING signaling pathway in inducing innate immune response has been firmly established (49, 50), its effect on the outcomes of inflammation-related diseases including cancer could be fundamentally distinct and even opposite. Recently, much effort has been put in reconciling the distinct impact of cGAS-STING activation on cancers by hypothesizing that while transient engagement of cGAS-STING results in acute inflammation and thereby leads to tumor control, sustained activation of this pathway and the consequent chronic inflammation may instead promote tumor progression (51, 52). In this context, multiple factor such as the cell types with cGAS-STING activation (e.g., tumor cells, accessory cells, and T cells), the activation/differentiation status of these cells, the source of STING agonists (e.g., mtDNA, nucDNA, and artificial chemical compounds), and cancer stages (precancerous, early and late) might affect the inflammation status and hence distinct disease outcomes. For example, in renal epithelial cells, mitochondrial fumarate accumulation caused by fumarate hydratase (FH) deficiency can lead to constant leakage of mtDNA into the cytoplasm, thereby activating the cGAS-STING-TBK1 signaling axis and triggering a continuous innate immune response. This mechanism has been proven to be closely related to the pathogenesis of hereditary leiomyomatosis and renal cell carcinoma (HLRCC) (53). In contrast, we found that DMF treatment is effective in treating CC through activating cGAS-STING-TBK1 pathway in mouse tumor models. Several explanations can be offered to reconcile this seemly discrepancy. First, cancer type and stages are different in these two studies (developed CC vs developing HLRCC); Second, DMF treatment of CC induces a transient activation of mtDNA-cGAS-STING pathway, while FH deficiency in renal epithelial cells results in constantly elevated mtDNA-cGAS-STING response, likely leading to acute and chronic inflammation, respectively. Therefore, activating cGAS-STING pathway by DMF treatment may represent a promising therapeutic drug for treating CC. Indeed, high expression of *STING* is positively correlated with the prognosis of human CC (Figure S3 F). Furthermore, as an FDA-approved drug with documented safety profiles, DMF functions through inducing the leakage of endogenous mtDNA into cytoplasm, distinguishing it from current exogenous STING agonists that are mostly artificial chemical compounds with safety profiles remained to be examined, thus making this therapeutic strategy even more attractive. However, given the aforementioned complexity regarding the role of cGAS-STING in cancer, cautions should be taken before translating it into clinical practice and extending it to other cancers.

The mechanism by which DMF promotes tumor regression resembles the immunogenic enhancement effect induced by chemotherapy or radiotherapy (54). Since combined strategies involving chemoradiotherapy and immunotherapy are currently undergoing clinical trials for various cancer types (55–58), the low-side effect profile of DMF thus make it a strong candidate for combinatorial therapies with current immunotherapies. Indeed, our findings demonstrate that DMF enhances the efficacy of PD-1 blockade and adoptive TIL therapy in mouse models. These synergic effects could be attributable to the following mechanisms. On the one hand, DMF treatment may increase the infiltration and cytotoxic differentiation of either PD-1 blockade-reinvigorated CTLs or adoptively transferred TILs, thus make both immunotherapies more effective. On the other hand, STING activation might upregulate PD-L1 expression in tumor cells (59), which could in turn increase tumors’ sensitivity to ICB therapy. Given the limited efficacy of ICB monotherapy in CC (with response rates of only 14.6%-27.8%) (6), and the diverse patient response to TIL therapy (ranging from no-responsive to complete tumor regression), the synergistic effect of DMF with ICB and TILs demonstrated in this study provides insight into developing novel effective combinatorial strategies in treating advanced CC.

Despite these promising prospects, several challenges remain. The effects of tumor heterogeneity, immune evasion mechanisms, and the potential systemic inflammation caused by prolonged STING activation on DMF treatment need to be carefully examined in further investigations. Moreover, future clinical trials are warranted to optimize DMF dosing regimens and examine the therapeutic efficacies of DMF monotherapy and its combination with TIL therapy or immune checkpoint inhibitors.

## Conclusions

Immune cell chemotaxis dysfunction in solid tumors remains a central problem to effective cancer immunotherapy. Our study discovered that DMF is capable of reprogramming CC cells to promote CD8^+^ T cell dependent anti-tumor immunity through activating the mtDNA-cGAS-STING pathway. DMF treatment induces mitochondrial dysfunction in CC cells, leading to the release of mtDNA into cytoplasm, which activates the cGAS-STING pathway and type I interferon response, and subsequently induces the secretion of CCL5 and CXCL10, thereby promoting CD8^+^ T cell recruitment. Importantly, in syngeneic mouse models, DMF demonstrates synergistic activity with PD-1 blockade, while in PDX models it significantly enhances TIL functionality. These findings provide viable strategies to improve the therapeutic efficacy of current immunotherapies.

## Acknowledgements

We thank Dr. Zhiyin Song for his assistance in mitochondrial imaging. We also extend our gratitude to all individuals and organizations that contributed to the research and preparation of this manuscript.

## Author contributions

Y.F. Huang, H. Wang and J.Y. Han contributed to the conception, design of the study. Y.F. Huang, H. Wang, T. Ye and J.Y. Han contributed to funding acquisition. H. Jiang and L.T. Liu contributed to the development of methodology. H. Jiang, L.T. Liu, S. He, S. Qu, Y.F. Yang, G.J. Kang Y.W. Zhang, Z.X. Wang and W.J. Tian contributed to the acquisition of data. M. Wu, H.Y. Liu, Y. Chen, L.M. Wang and Q.Q. Wang contributed to the interpretation of data.

## Ethics approval and consent to participate

This study was reviewed and approved by the Institutional Animal Care and Use Committee (IACUC) of Huazhong University of Science and Technology. ([2024] IACUC Number:4334).

---

**Fig S1.**
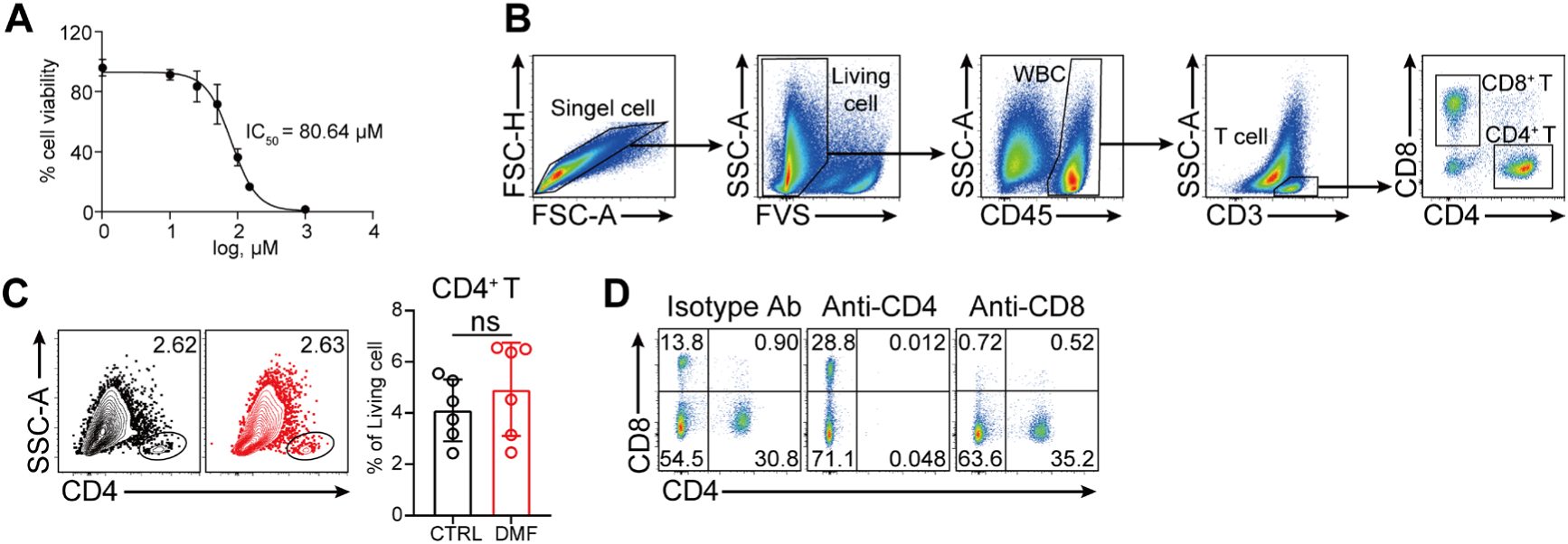
**A**, Survival curves of TC-1 cells treated with different concentrations of DMF. **B**, Flow cytometry analysis workflow for tumor tissues from mice. **C**, Percentage of infiltrating CD8⁺ T cells in tumor of immunocompetent mice. (n = 6). **D**, Proportion of CD4⁺ or CD8⁺ T cells in the spleens of mice after treatment with CD4/CD8 depleting antibodies. Data are presented as the mean ± SD. *P* values were calculated using unpaired two-tailed Student’s *t* test (**C**), ns indicating no significant difference.

**Fig S2.**
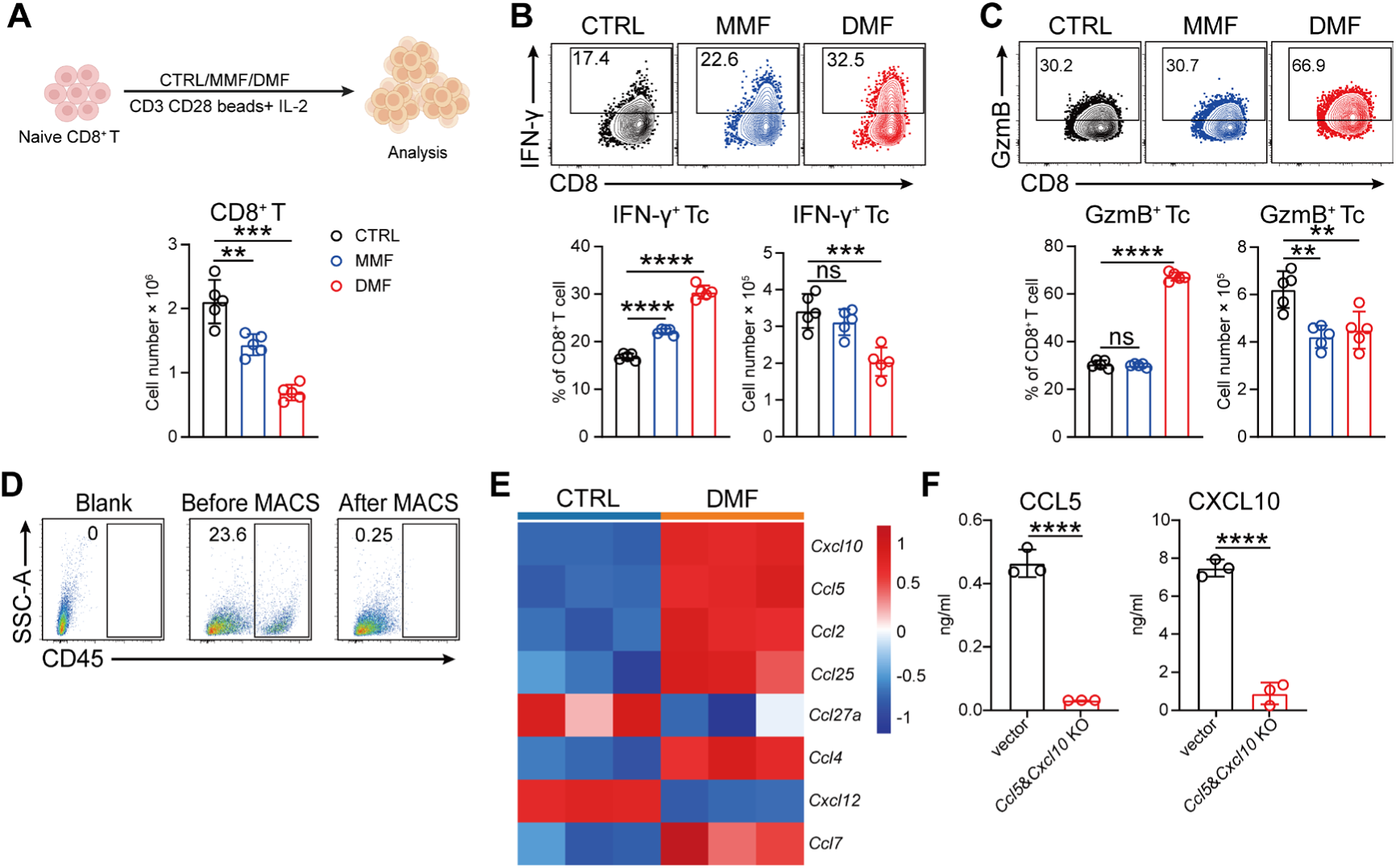
**A**-**C**, Naïve CD8^+^ T cells were treated with 100 μM MMF or 50 μM DMF, and simultaneously cultured with CD3&CD28 beads and IL-2 in lymphocyte culture medium for 72 h. The number of T cell (**A**), the number and percentage of IFN-γ^+^ CD8^+^ T cell (**B**), and GzmB^+^ CD8^+^ T cell (**C**) were measured. **D**, Purity of CD45^-^ tumor cells in tumor tissue before and after magnetic bead sorting wase analyzed by flow cytometry. **E**, Chemokine gene expression in TC-1 cells treated with or without 50 μM DMF. **F**, Concentrations of CXCL10 and CCL5 in the supernatant of TC-1 vector or CCL5 & CXCL10 knockout cells. Data are the mean ± SD. *P* values were calculated using unpaired two-tailed Student’s *t* test (**F**), one-way ANOVA for Dunnett’s multiple comparisons test (**A**-**C**). \*\**P* < 0.01, \*\*\**P* < 0.001, \*\*\*\**P* < 0.0001, and ns indicating no significant difference.

**Fig S3.**
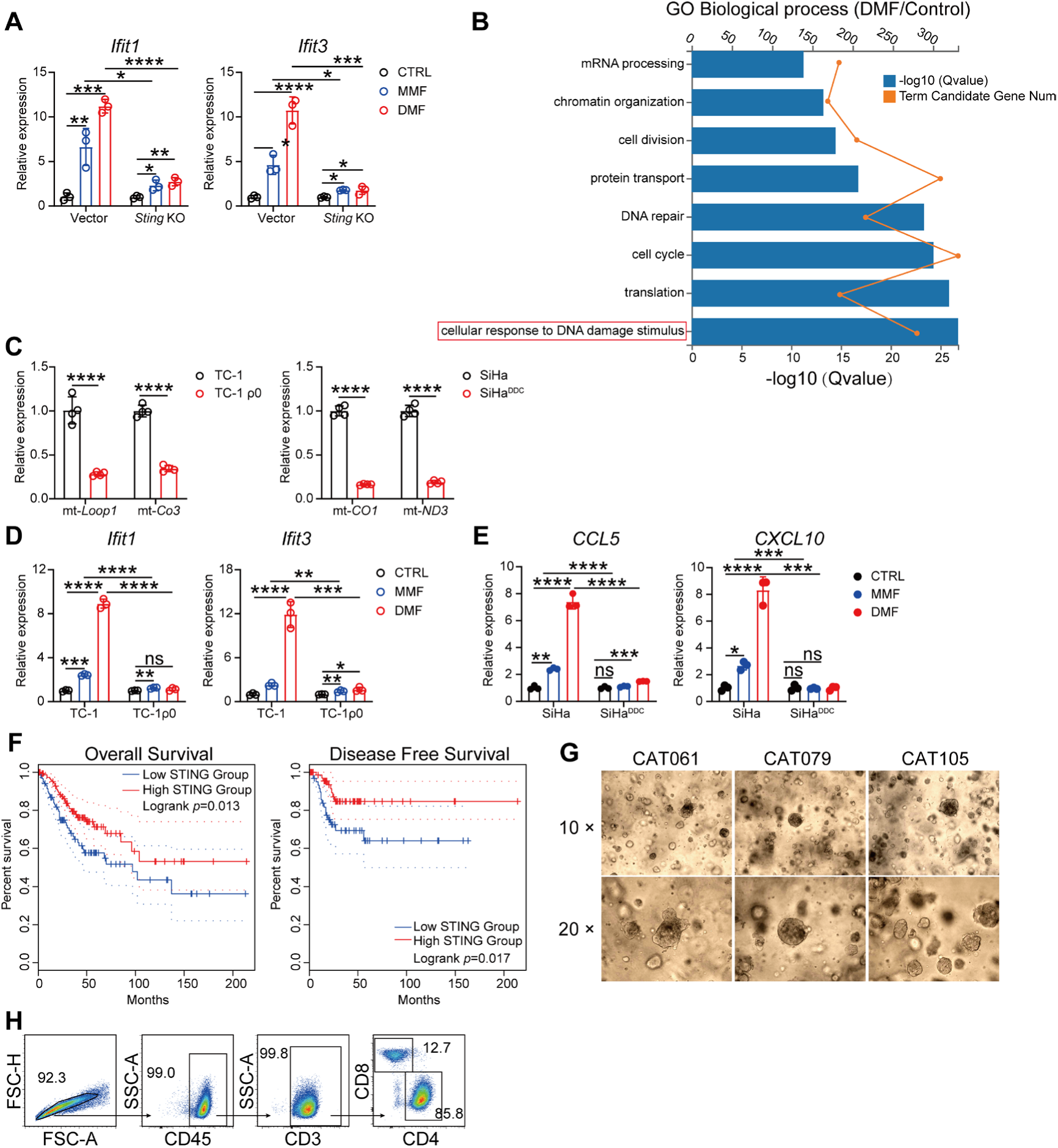
A, Transcriptional levels of *Ifit1* and *Ifit3* were quantified by qRT-PCR in vector or *Sting* KO TC-1 cells treated with 100 μM MMF or 50 μM DMF for 24 h. **B**, GO enrichment analysis of TC-1 cells treated with or without 50 μM DMF. **C**, mtDNA content was assessed by qRT-PCR following 20 days of treatment with 100 ng/ml EthBr in TC-1 cells or 100 nM DDC in SiHa cells. **D**, Transcriptional levels of *Ifit1* and *Ifit3* were quantified by qRT-PCR in TC-1 and TC-1ρ0 cells treated with 100 μM MMF or 50 μM DMF for 24 h. **E**, Transcriptional levels of *CCL5* and *CXCL10* were quantified by qRT-PCR in SiHa and SiHa^DDC^ cells treated with 100 μM MMF or 50 μM DMF for 24 h. **F**, Kaplan-Meier analysis comparing overall survival and disease-free survival in CC patients with low versus high *STING* gene expression. Analysed with http://gepia2.cancer-pku.cn/. **G**, Representative images of organoids derived from cervical cancer patients, captured under 10 × and 20 × magnification. (n = 3 patients). **H**, TIL subpopulations expanded ex vivo from cervical cancer patient was analyzed by flow cytometry. Data are presented as the mean ± SD. *P* values were calculated using two-way ANOVA for Tukey’s multiple comparisons test (**A**, **C**-**E**), with significance levels defined as \**P* < 0.05, \*\**P* < 0.01, \*\*\**P* < 0.001, \*\*\*\**P* < 0.0001, and ns indicating no significant difference.

**Table 1.**
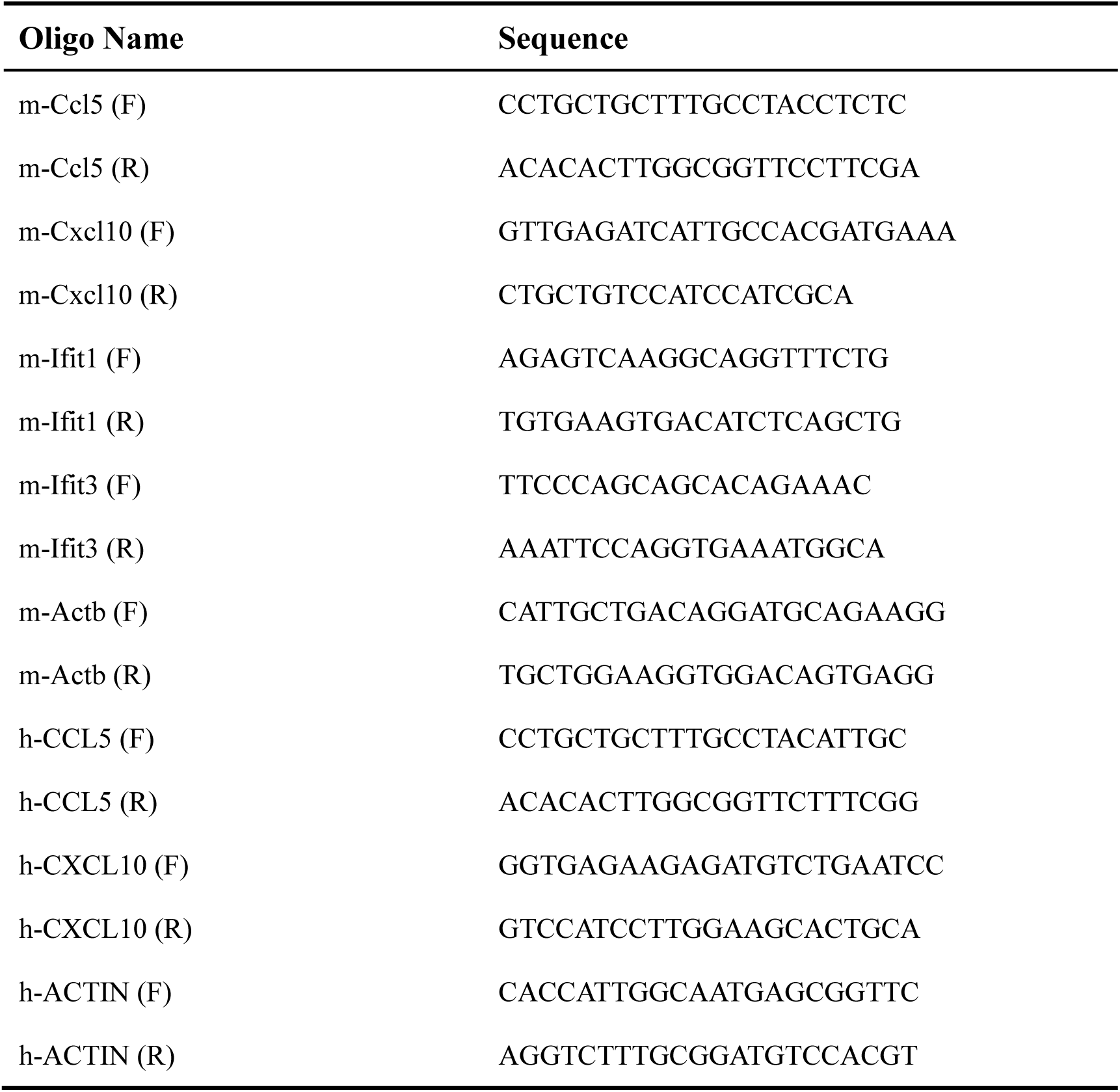
qRT-PCR primer.

**Table 2.**
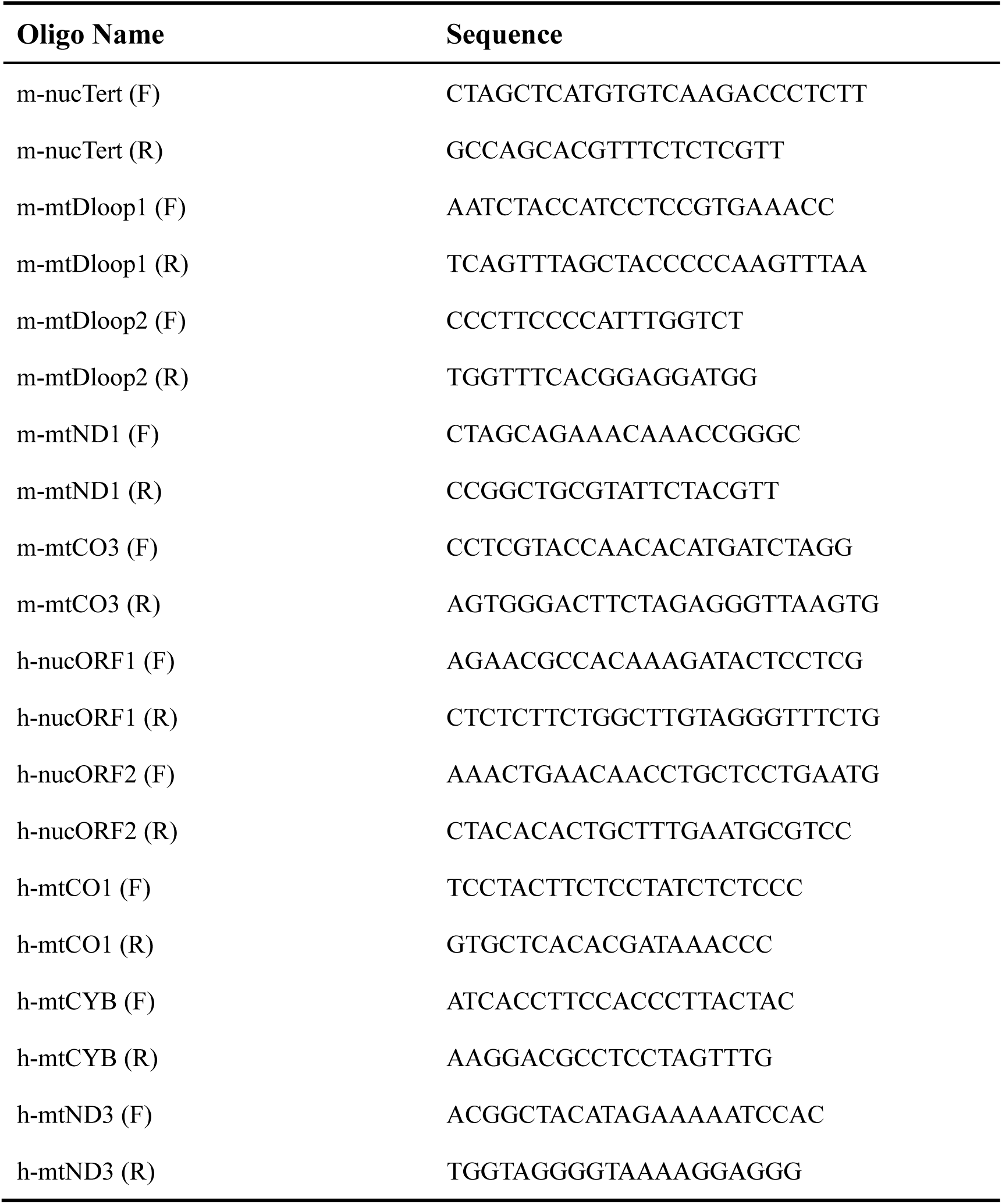
DNA assay.

**Table 3.** Lymphocyte culture medium.

| **Product** | **Concentration** | **Catalog** |
| --- | --- | --- |
| RPMI 1640 | / | Gibco (10379144) |
| FBS | 10 % | Gibco (A4766801) |
| GlutaMax | 1X | Gibco (35050087) |
| HEPES | 10 mM | Gibco (15630-106) |
| Penicillin/Streptomycin | 1 mg/ml | Beyotime (C0222) |
| IL-2 | 10 U/ml | Peprotech (AF-212-12) |
| β-mercaptoethanol | 50 μM | Sigma (444203) |

**Table 4.** PDXO culture med.

| **Product** | **Concentration** | **Catalog** |
| --- | --- | --- |
| Advanced DMEM/F12 | / | Gibco (12634010) |
| GlutaMax | 1X | Gibco (35050087) |
| HEPES | 10 mM | Gibco (15630-106) |
| Penicillin/Streptomycin | 1 mg/ml | Beyotime (C0222) |
| B27 | 1X | Gibco (17504044) |
| Human EGF | 10 ng/ml | Peprotech (AF-100-15) |
| Human Noggin | 100 ng/ml | Peprotech (120-10C) |
| Human FGF-7 | 25 ng/ml | Peprotech (100-19) |
| N-acetyl-L-cysteine<sup>9</sup> (NALC) | 1.25 mM | MCE (HY-B0215) |
| Nicotinamide (NCTA) | 2.5 mM | MCE (HY-B0150) |
| TGF-β I Receptor inhibitor (A83-01) | 500 nM | MCE (HY-10432) |
| ROCK inhibitor (Y-27632) | 10 μM | MCE (HY-10071) |
| Forskolin | 10 μM | MCE (HY-15371) |
| P38 MAPK inhibitor (SB 202190) | 1 μM | MCE (HY-10295) |

## Notes

### Competing Interest Statement

The authors have declared no competing interest.

### Funding

- National Natural Science Foundation of China: 82273211, 82373260, 82403864
- Nature Science Foundation of Hubei Province: 2021CFB346, 2022CFB231
- National Key Research and Development Program of China: 2021YFC2701204
- Key R&D Program of Zhejiang: 2022C03013

